# Characterization and structural prediction of the putative ORF10 protein in SARS-CoV-2

**DOI:** 10.1101/2020.10.26.355784

**Authors:** Noah A. Schuster

**Affiliations:** Department of Biology, DePauw University, Greencastle, IN

**Keywords:** Coronavirus, SARS-CoV-2, ORF10, Accessory Protein, Structure Prediction

## Abstract

Upstream of the 3’-untranslated region in the SARS-CoV-2 genome is ORF10 which has been proposed to encode for the ORF10 protein. Current research is still unclear on whether this protein is synthesized, but further investigations are still warranted. Herein, this study uses multiple bioinformatic tools to biochemically and functionally characterize the ORF10 protein, along with predicting its tertiary structure. Results indicate a highly ordered, hydrophobic, and thermally stable protein that contains at least one transmembrane region. This protein also possesses high residue protein-binding propensity, primarily in the N-terminal half. An assessment of forty-one missense mutations reveal slight changes in residue flexibility, mainly in the C-terminal half. However, these same mutations do not inflict significant changes on protein stability and other biochemical features. The predicted model suggests the ORF10 protein contains a β-α-β motif with a β-molecular recognition feature occurring in the first β-strand. Functionally, the ORF10 protein could be a membrane protein. A single pocket was identified in this protein but found to possess low druggability. The ORF10 itself consists of two distinct lineages: the SARS-CoV lineage and the SARS-CoV-2 lineage. Evidence of strong positive selection (dN/dS = 4.01) and purifying selection (dN/dS = 0.713) were found within the SARS-CoV-2 lineage and SARS-CoV lineage, respectively. Collectively, these results continue to assess the biological relevance of ORF10 and its putatively encoded protein, thereby aiding in diagnostic and possibly vaccine development.

## 1. Introduction

An outbreak of coronavirus disease 2019 (COVID-19) in Wuhan, China has resulted in a global pandemic causing, at the time this manuscript was prepared, well over 102,135,000 cases and 2,210,000 deaths.^1^ The virus responsible for causing COVID-19, severe acute respiratory syndrome coronavirus type 2 (SARS-CoV-2), contains an individual copy of a single-stranded positive-sense RNA genome that is ∼29.8 kilobases in length.^2^ In particular, the 3’-terminus encompasses multiple open reading frames (ORFs) that encode four main structural proteins: the spike (S), membrane (M), envelope (E), and nucleocapsid (N) proteins.^2^ In addition, shorter length ORFs have been detected and proposed to encode for approximately nine different accessory proteins (3a, 3b, 6, 7a, 7b, 8, 9b, 9c, 10) in the SARS-CoV-2 genome.^2,3^ Past research indicates these accessory proteins are dispensable for viral growth *in vitro*; however, the genes encoding for these proteins are maintained in the coronavirus (CoV) genome, suggesting they might play important roles within the environment of the infected host.^4^

To some degree, all of these accessory proteins have been structurally and/or functionally characterized. For example, the 3a protein was shown to induce apoptosis whereas the 9b and 9c proteins were suggested to be involved in membrane interactions during virion assembly and host-virus interactions, respectively.^5,6^ Prior work has been able to detect for the expression of RNA transcripts corresponding to these proteins in SARS-CoV-2; however, RNA transcripts belonging to ORF10 have not been observed in large amounts.^3,7^ Pancer *et al*. 2020 had also identified several SARS-CoV-2 variants with prematurely terminated ORF10 sequences, finding that disease was not attenuated, transmission was not hindered, and replication proceeded similarly to strains possessing intact ORF10 sequences.^8^ In light of these findings, there is some uncertainty regarding the biological relevance of ORF10 in SARS-CoV-2, with suggestions that its genome annotation should be amended. As an unintended consequence, this putatively encoded protein within SARS-CoV-2 has remained somewhat understudied.

Found upstream of the 3’-untranslated region (3’-UTR), ORF10 (117 nt long) encodes for a protein that is 38 amino acids in length.^9^ Although there exists sequences homologous to the ORF10 protein in other closely related bat and pangolin CoVs, there are no experimentally derived crystallographic structures for the ORF10 protein.^10,11^ Studies attempting to predict the secondary structural elements of this protein indicate one α-helix and, depending on the study, two β-strands.^12,13^ The ORF10 protein also lacks significant levels of disorder; however, a short molecular recognition feature (MoRF) likely spans residues 3-7.^12^ This protein is hydrophobic with the α-helix having been identified as a possible transmembrane (TM) helix as well.^14^

The exact implications are unclear, but the ORF10 protein contains high numbers of promiscuous cytotoxic T lymphocyte (CTL) epitopes, primarily on the α-helix.^12,15^ A recent study by Liu *et al*. 2020 found that in severe cases of COVID-19, the ORF10 was overexpressed when compared against ORF10 expression levels in moderate cases.^16^ If the ORF10 protein is synthesized, this may help to partially explain the nature behind severe cases of COVID-19. The large number of CTL epitopes in conjunction with the overproduction of the ORF10 protein could result in elevated immune responses towards SARS-CoV-2, leading to some potentially fatal immunopathological outcomes.

The ORF10 protein was shown to associate with the CUL2^ZYG11B^ complex by interacting with the substrate adapter ZYG11B, thereby overtaking the complex to modulate aspects of ubiquitination and enhance viral pathogenesis.^13^ However, the mechanisms and residues involved in this interaction have not been described in detail. Alternatively, ORF10 may act itself or serve as a precursor for other RNAs in regulating gene expression/replication, translation efficiency, or interfering with cellular antiviral pathways.^7^

Two lineages for the ORF10 have been observed. The SARS-CoV lineage consists of sequences interrupted by a stop codon (TAA) encoding for a truncated protein and was found to be undergoing neutral evolution (dN/dS = 1.18).^14^ The SARS-CoV-2 lineage consists entirely of uninterrupted sequences encoding for a complete protein and was found to be undergoing strong positive selection (dN/dS = 3.82) without relaxed constraints.^14^ This has led to the possibility that the ORF10 in SARS-CoV-2 and closely related CoVs might encode a conserved and functional protein. Furthermore, the ORF10 present in SARS-CoV-2 and closely related CoVs is thought to have evolved from a mutation occurring within the ancestral stop codon originally found in the SARS-CoV lineage at nucleotide position 76.^17^

Collectively, there still remains few efforts spent on investigating the ORF10 protein within SARS-CoV-2. Herein, this current study uses a bioinformatic approach to characterize this protein, along with providing a more updated phylogenetic and evolutionary analysis. This study also examines the structural and biochemical effects of mutations found in this protein, along with assessing its potential as an antiviral drug target.

## 2. Materials and Methods

### 2.1 Sequence, Phylogenetic, and Evolutionary Analysis

The nucleotide (NC_045512.2) and protein (YP_009725255.1) reference sequences for ORF10 in SARS-CoV-2 were acquired from the NCBI RefSeq Database. BLASTn was used to collect sequences from both distantly and closely related CoVs (Table S1).^18^ Sequence alignments were performed using MUSCLE on the MEGA-X v10.1.7 software.^19,20^ Alignment reliability and total similarity were determined by overall mean distance and calculated using the *p-distance* substitution model. If the p-distance was found to be ≤ 0.7 then the alignment was considered reliable and suitable for subsequent analyses.^21^

A single phylogenetic gene tree was constructed using the neighbor-joining method and visualized on MEGA-X.^22^ Uniform rates among sites and complete deletion were used. The phylogeny was tested using the bootstrap method.

The average nonsynonymous/synonymous substitution (dN/dS) rate was calculated using the single-likelihood ancestor counting (SLAC) method.^23^ To determine either relaxed (k < 1) or intensified (k > 1) selection, the RELAX method was utilized.^24^ Both methods were performed using the datamonkey webtool (http://datamonkey.org/). Wu-Kabat variability coefficients were calculated using the protein variability server (http://imed.med.ucm.es/PVS/).^25^

### 2.2 Protein Characterization and Secondary Structure Prediction

Per-residue disorder scores were determined using several online tools as previously described.^26^ Scores were then used to calculate average protein disorder. To detect phosphorylation sites, the DEPP server (http://www.pondr.com/cgi-bin/depp.cgi) was used. A hydrophobicity plot was generated using ProtScale.^27^ The grand average of hydropathicity (GRAVY), aliphatic index, and instability index were determined using ProtParam.^27^ The propensity of residues involved in protein-binding were evaluated by SCRIBER.^28^ Secondary structural elements were predicted using PSIPRED v4.0.^29^ TM regions were predicted using TMPred.^30^

### 2.3 Protein Modeling and Evaluation

The webserver IntFOLD was employed to make use of an *ab initio* modeling approach in constructing the ORF10 protein.^31^ Models were evaluated based on quality and confidence scoring. The best model was then refined using the 3Drefine webserver.^32^ Of the models generated post-refinement, the one with a higher qualitative model energy analysis (QMEAN) global Z-score was viewed as most favorable.^33^ This final model was evaluated by PROCHECK and ERRAT on the SAVES v6.0 webtool (https://saves.mbi.ucla.edu/).^34-36^ This theoretical structure was deposited in ModelArchive (https://modelarchive.org/) and given the ID: ma-9yzbf. Global and local quality comparisons against two previously constructed models were also performed.^37,38^

The ProFunc server predicted the function of the ORF10 protein based on the newly produced model.^39^ The online web server DoGSiteScorer was used to predict and describe pockets within the ORF10 protein.^40^ A druggability score was returned between 0 and 1. The higher the score, the more druggable the pocket was estimated to be. UCSF Chimera v1.15 was used to visualize and obtain high-quality images of protein models.^41^ Electrostatic surfaces were generated based on the AMBER ff14SB charge model. Hydrophobicity surfaces were produced according to the standard Kyte-Doolittle scale.

### 2.4 Mutation Analysis

Using forty-one ORF10 protein variants, the effect these mutations had on protein stability and other biochemical features were determined. The same methods that were used to determine the GRAVY, instability index, aliphatic index, and per-residue disorder scores were used to evaluate the effects of these mutations. The effect each mutation had on protein stability was determined using the I-MUTANT v2.0 webtool.^42^ The free energy change value (DDG) predictor was used to assess changes in stability brought on by each mutation. If a negative value is returned, then the stability of the protein decreases. If a positive value is returned, then the stability of the protein increases. The designated temperature and pH used were 25°C and 7, respectively.

## 3. Results

### 3.1 Sequence, Phylogenetic, and Evolutionary Analysis

The whole nucleotide alignment (p-distance = 0.08) had 92% similarity. The phylogenetic gene tree revealed two lineages (Figure 1a). The SARS-CoV lineage (p-distance = 0.04) had 96% similarity whereas the SARS-CoV-2 lineage (p-distance = 0.08) had 92% similarity. An alignment of protein sequences taken from the SARS-CoV-2 lineage revealed a total of thirty-one conserved sites (Figure 1b). A Wu-Kabat plot displayed higher variability coefficients in the C-terminal half (20-38) rather than the N-terminal half (1-19) of the ORF10 protein, with amino acid position twenty-five presenting the highest variability coefficient in the lineage (Figure 1c).

**Figure 1.**
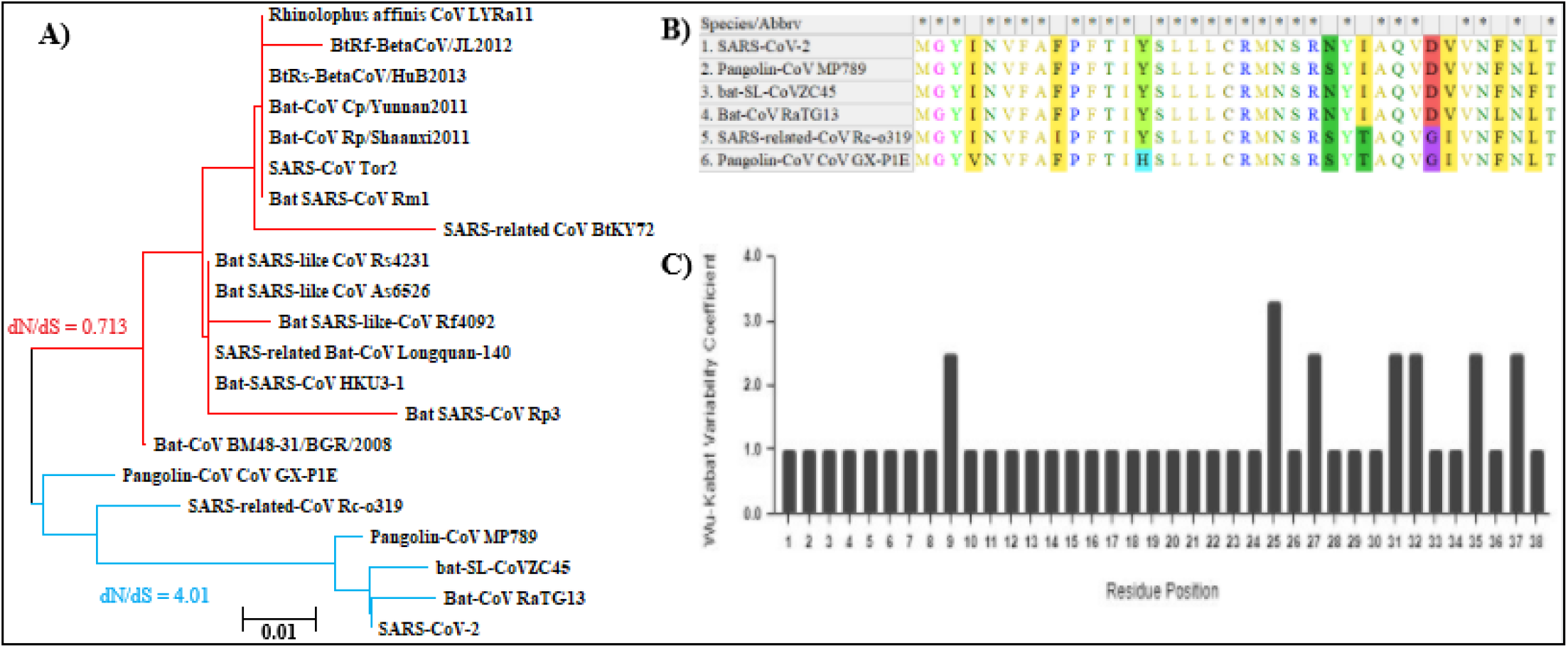
**(A)** Phylogenetic analysis of ORF10 sequences. The SARS-CoV lineage has branches shown in red whereas the SARS-CoV-2 lineage has branches shown in blue. Their corresponding dN/dS ratios calculated by SLAC are also shown. The phylogeny was inferred using the Jukes-Cantor method. The sum of branch lengths was 0.192 and the tree was produced out of 1,000 replicates. The bootstrap values are not shown. **(B)** Screen capture of the MUSCLE alignment for ORF10 proteins in the SARS-CoV-2 lineage. Conserved positions are indicated with an asterisk. **(C)** Wu-Kabat plot highlighting amino acid variations found in the ORF10 protein taken from the SARS-CoV-2 lineage. The consensus sequence was utilized as the reference sequence in generating this plot.

The SARS-CoV lineage had a dN/dS = 0.713 to indicate purifying selection. Oppositely, the SARS-CoV-2 lineage had a dN/dS = 4.01 (Figure 1a). To determine if this dN/dS ratio was either the result of positive or relaxed purifying selection, the RELAX method was used. In doing so, a k = 6.79 was obtained; therefore, the hypothesis of a relaxed constraint was rejected in favor of strictly positive selection acting upon the SARS-CoV-2 lineage.

### 3.2 Protein Characterization and Secondary Structure Prediction

A per-residue disorder plot for the ORF10 protein in SARS-CoV-2 showed that residues within both termini held the highest disorder scores whereas moving inwards resulted in gradually decreasing scores (Figure 2). The average disorder score for the ORF10 protein was 0.095, which indicates a highly ordered protein. No specific phosphorylation sites had been detected within this protein (Table S2). The GRAVY score was 0.637 to indicate a very hydrophobic protein. The aliphatic and instability indices were 107.03 and 16.06, respectively. Both of these values suggest a thermally stable protein. The hydrophobicity plot uncovered two hydrophobic regions covering residues 3-19 and 28-36, along with a hydrophilic region covering residues 20-27 (Figure 3a). As for protein-binding propensity, residues in the N-terminal half had a tendency to possess greater scores than residues found in the C-terminal half (Figure 3b). In particular, residues 1-14 presented with a stretch of relatively high propensity scores to implicate this region’s possible involvement in protein-binding (Figure 3b).

**Figure 2.**
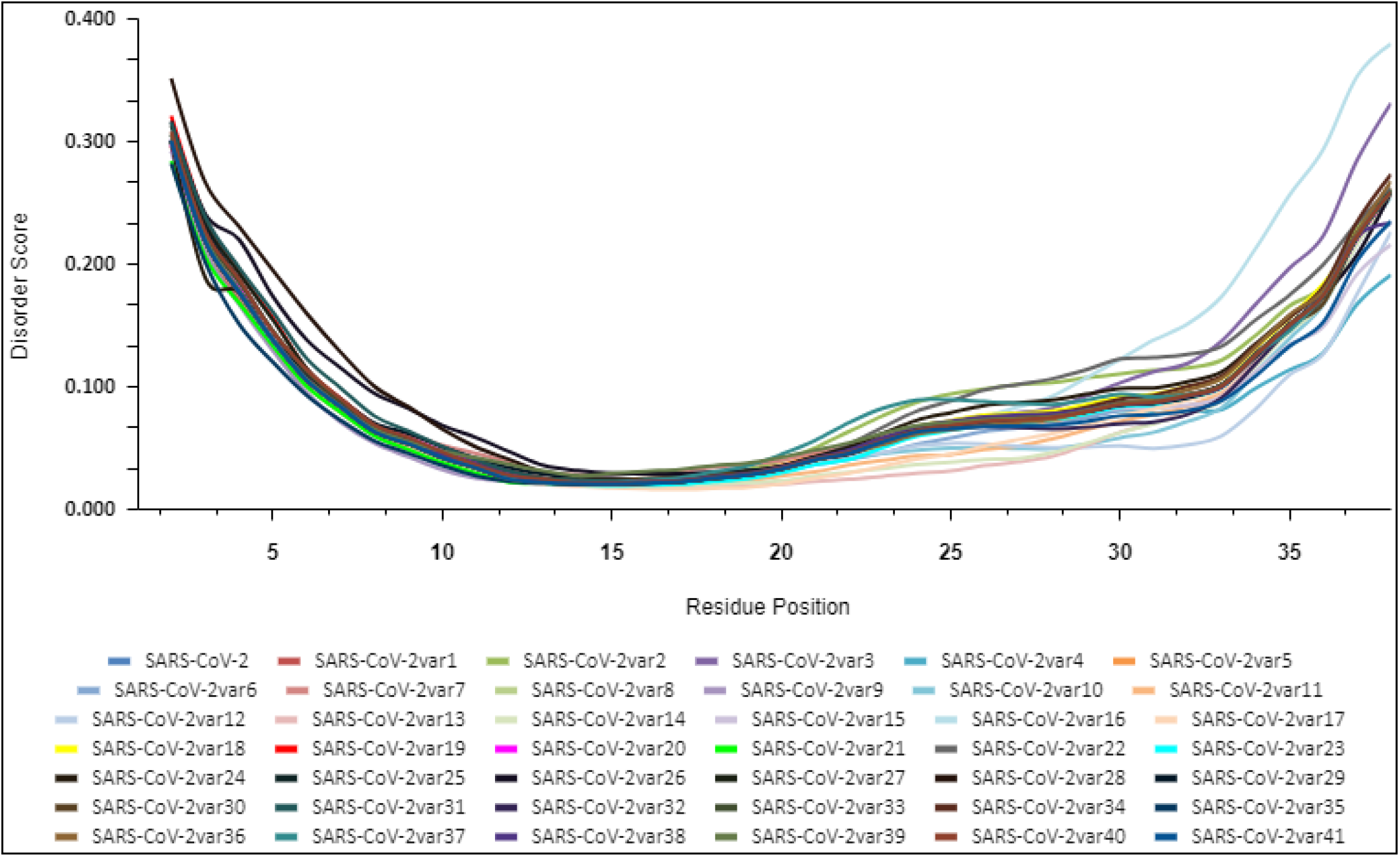
Per-residue disorder plot for the ORF10 protein in SARS-CoV-2, along with twenty-two variants. Scores ≥ 0.5 indicate disorder whereas scores between 0.25 and 0.5 indicate highly flexible residues. Any score that exists between 0.1 and 0.25 suggests moderate flexibility. Scores ≤ 0.1 indicate rigidity.

**Figure 3.**
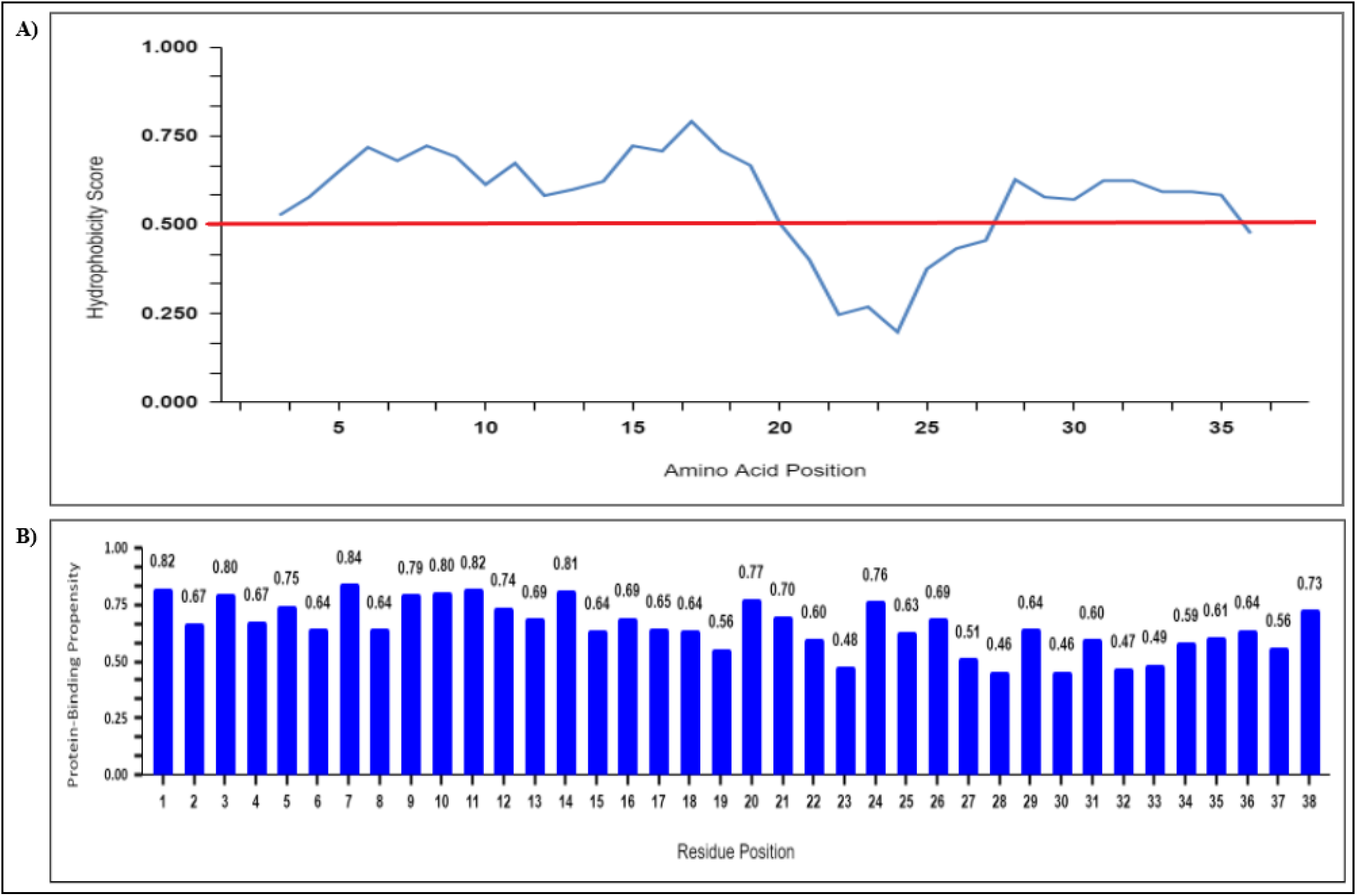
**(A)** Hydrophobicity plot for the SARS-CoV-2 ORF10 protein. In generating this plot, scores were normalized on a 0-1 scale and a 5-residue window was used. The red line displays the threshold where values above this line indicate hydrophobic regions and values below this line indicate hydrophilic regions. Residues 1-2 and 37-38 are not shown on this plot. **(B)** Protein-binding propensity scores for amino acids within the SARS-CoV-2 ORF10 protein. The value of each score is shown above its corresponding box. As a general rule, a higher score reflects an increased likelihood that a specific residue is going to engage in a protein-binding interaction.

The prediction of secondary structures indicates that a single α-helix (residues 11-21) and two β-strands (residues 4-8 & 26-34) occur in the ORF10 protein (Table S3). Furthermore, only one TM region was predicted to span residues 3-19, corresponding to one of the hydrophobic regions that was shown in the hydrophobicity plot (Figure 3a).

### 3.3 Protein Modeling and Predictive Function

IntFOLD produced five different ORF10 protein models (Table S4). The model having the lowest p-value (3.33e-03) and highest quality score (0.3714) was subjected to refinement. Of the five protein models generated post-refinement, the model with a QMEAN Z-score of −0.66 was selected as most favorable (Table S5). This final model had an ERRAT score of 93.333 and a Ramachandran plot indicated that 94% of residues (32 total) existed in the most favorable region while only 6% of residues (2 total) existed in the additional allowed region (Figure S1). Functional analysis results by ProFunc have been summarized (Table 1). The predicted ontological terms were based on “long shot” hits using a reverse template comparison against known Protein Data Bank structures; therefore, these functional characteristics should not be viewed as definitive.

**Table 1.**
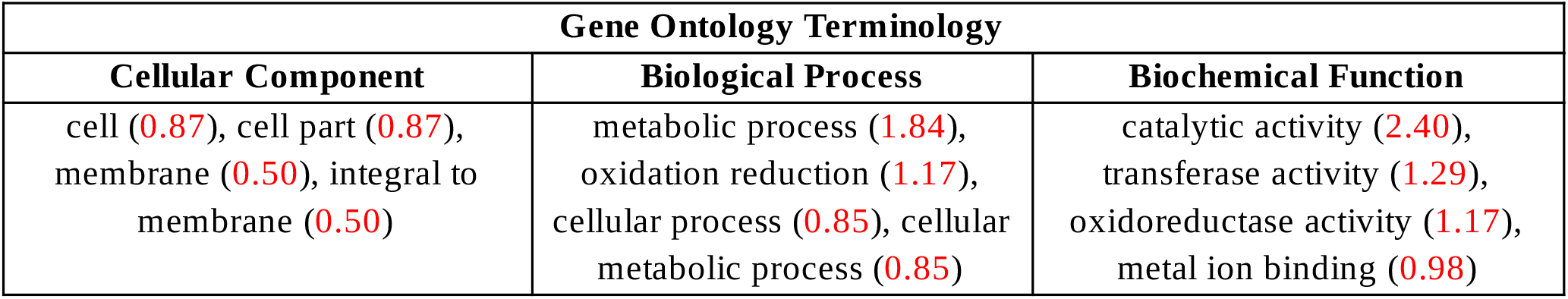
Predicted gene ontology terms for the ORF10 protein in SARS-CoV-2. These terms are the most commonly appearing terms found in the hits obtained by using the ProFunc analysis method. Each of these terms’ scores are based on the number of times they independently occur during the analysis and are shown in red and within parentheses.

Structurally, the model presented with a β-α-β motif spanning residues 3-31, along with a 3/10-helix spanning residues 34-37 (Figure 4a). An electrostatic surface map revealed two regions of positive charge and one region of negative charge (Figure 4c). Electropositive regions were influenced by R20 and R24 whereas the electronegative region was influenced by D31. As expected, a majority of the protein’s surface was hydrophobic; however, hydrophilic regions did appear (Figure 4d). For example, residues spanning 20-27 presented with surface coloring that reflected hydrophilic character, as was shown in the hydrophobicity plot (Figure 3a).

**Figure 4.**
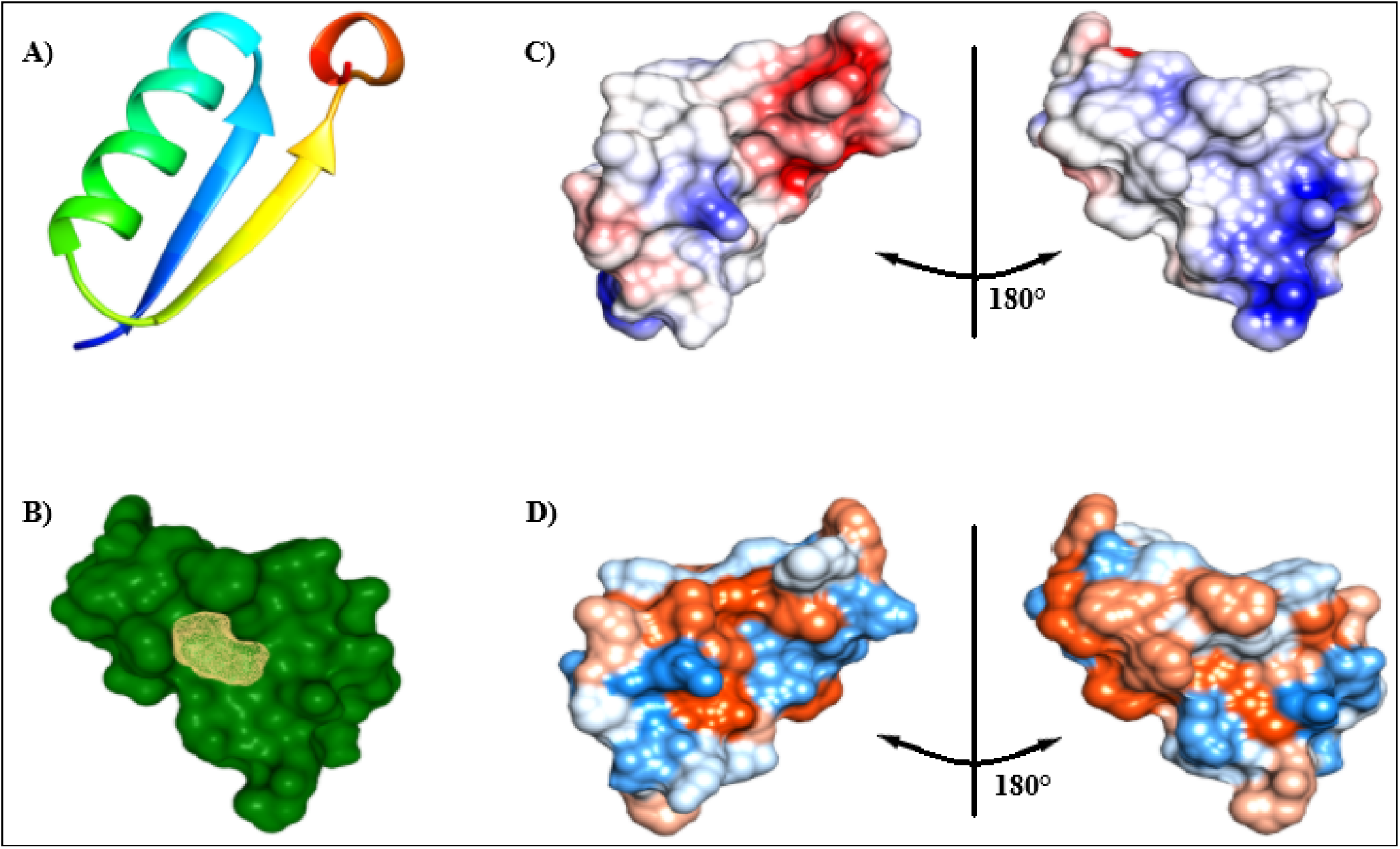
**(A)** Ribbon diagram of the ORF10 protein in SARS-CoV-2. **(B)** Surface of the ORF10 protein (green), along with the location of the pocket identified by DoGSiteScorer (yellow). **(C)** Electrostatic surface map of the ORF10 protein. Red indicates regions having negative charge whereas blue indicates regions having positive charge. White indicates regions having neutral charge. **(D)** Hydrophobicity surface map of the ORF10 protein. Blue indicates regions of hydrophilic character and orange-red indicates regions of hydrophobic character. White indicates regions of amphipathic character.

A single hydrophobic pocket was detected using DoGSiteScorer (Figure 4b). This pocket had a volume of 78.14 Å^3^, a surface area of 221.79 Å^2^, and a depth of 5.31 Å. This pocket had a total of 19 hydrophobic interactions, followed by 8 hydrogen bond acceptors and 3 hydrogen bond donors. The druggability score for this pocket was 0.14 and interpreted as a poor drug target.

Prior to this study, two ORF10 protein models were constructed. These models were built using either the QUARK or I-TASSER web server.^36,37^ Both models varied in placement of secondary structural elements and topology (Figure S2). The QMEAN global Z-scores for both models (QUARK: −3.88 / I-TASSER: −2.63) were lower, indicating the model produced in this study had higher global, and also local, quality scoring (Figure S3).

### 3.4 Mutation Analysis

One missense mutation per variant was observed. The longest conserved region only spanned residues 15-16 (Figure 5). Per-residue disorder scores in the N-terminus did not fluctuate as much when compared to per-residue disorder scores in the C-terminus (Figure 2). Regardless, the average disorder for each variant indicated highly ordered proteins. The predicted effects that each missense mutation had on protein structure, stability, and other biochemical features have been summarized (Table 2).

**Table 2.** Forty-one ORF10 protein variants from SARS-CoV-2, along with their location, corresponding mutations, and predicted structural and biochemical effects. NP - Nonpolar; P - Polar.

| SARS-CoV-2<br>ORF10 Variant | Nucleotide<br>Accession | Protein<br>Accession | Geographic<br>Location | Amino Acid<br>Change | Change in<br>Polarity | Change in<br>Stability | GRAVY<br>Score | Average<br>Disorder | Instability<br>Index | Aliphatic<br>Index |
| --- | --- | --- | --- | --- | --- | --- | --- | --- | --- | --- |
| var1 | MT259226.1 | QIS29991.1 | China | V6I | NP → NP | -1.14 | 0.645 | 0.094 | 27.62 | 110.26 |
| var2 | MT632596.1 | QKU54102.1 | USA | Y26H | NP → P | -1.68 | 0.587 | 0.105 | 27.62 | 107.63 |
| var3 | MT632956.1 | QKV08176.1 | USA | L37P | NP → NP | -2.25 | 0.495 | 0.108 | 15.30 | 97.37 |
| var4 | MT641656.1 | QKV37245.1 | Australia | T38I | P → NP | -1.25 | 0.774 | 0.086 | 16.06 | 117.89 |
| var5 | MT679202.1 | QLA48060.1 | USA | R20K | P → P | -2.39 | 0.653 | 0.095 | 24.64 | 107.63 |
| var6 | MT745708.1 | QLG76514.1 | Australia | N22T | P → P | -1.34 | 0.711 | 0.093 | 15.30 | 107.63 |
| var7 | MT750342.1 | QLG99793.1 | USA | I3M | NP → NP | -0.17 | 0.568 | 0.098 | 22.29 | 97.37 |
| var8 | MT764166.1 | QLI33453.1 | USA | Y26C | NP → P | 0.41 | 0.737 | 0.094 | 16.06 | 107.63 |
| var9 | MT252755.1 | QLJ57416.1 | USA | A8V | NP → NP | -0.01 | 0.700 | 0.092 | 16.06 | 112.63 |
| var10 | MT806775.1 | QLY88596.1 | USA | A28V | NP → NP | -0.54 | 0.700 | 0.087 | 13.83 | 112.63 |
| var11 | MT831635.1 | QMT54534.1 | USA | R24L | P → NP | -1.40 | 0.855 | 0.089 | 7.74 | 117.89 |
| var12 | MT834229.1 | QMT94417.1 | USA | D31Y | P → NP | -1.48 | 0.695 | 0.081 | 18.03 | 107.63 |
| var13 | MT834697.1 | QMT97141.1 | USA | S23F | P → NP | 0.29 | 0.732 | 0.085 | 7.04 | 107.63 |
| var14 | MT846564.1 | QMU93213.1 | USA | R24C | P → P | -1.66 | 0.821 | 0.086 | 16.32 | 107.63 |
| var15 | MT876527.1 | QNA70543.1 | Bangladesh | L37F | NP → NP | -0.09 | 0.611 | 0.090 | 12.11 | 97.37 |
| var16 | MT876555.1 | QNB17780.1 | Bangladesh | F35S | NP → P | -3.45 | 0.542 | 0.121 | 20.02 | 107.63 |
| var17 | MT877418.1 | QNC04532.1 | USA | R20I | P → NP | -0.31 | 0.874 | 0.089 | 16.06 | 117.89 |
| var18 | MT879619.1 | QNC49349.1 | Pakistan | V30L | NP → NP | -1.31 | 0.626 | 0.097 | 22.00 | 110.26 |
| var19 | MT892994.1 | QNG41574.1 | USA | I4L | NP → NP | 0.52 | 0.618 | 0.098 | 16.06 | 107.63 |
| var20 | MT893673.1 | QNG42985.1 | USA | P10S | NP → P | -2.23 | 0.658 | 0.096 | 5.92 | 107.63 |
| var21 | MT911797.1 | QNI23218.1 | USA | G2D | NP → P | -0.71 | 0.555 | 0.093 | 18.29 | 107.63 |
| var22 | MT913142.1 | QNI25281.1 | USA | V30A | NP → NP | -2.55 | 0.574 | 0.106 | 19.77 | 102.63 |
| var23 | MW446204.1 | QQL06028.1 | USA | L17F | NP → NP | -0.12 | 0.611 | 0.094 | 16.06 | 97.37 |
| var24 | MW446223.1 | QQL06256.1 | USA | I27V | NP → NP | -1.07 | 0.629 | 0.099 | 16.06 | 105.00 |
| var25 | MW446840.1 | QQL12865.1 | USA | N22D | P → P | -1.19 | 0.637 | 0.095 | 21.13 | 107.63 |
| var26 | MW433744.1 | QQK54029.1 | USA | F9S | NP → P | -3.38 | 0.542 | 0.105 | 22.56 | 107.63 |
| var27 | MW425708.1 | QQJ94693.1 | USA | I4V | NP → NP | -0.63 | 0.629 | 0.097 | 16.06 | 105.00 |
| var28 | MW405472.1 | QQE14148.1 | USA | F7S | NP → P | -3.35 | 0.542 | 0.107 | 16.06 | 107.63 |
| var29 | MW405483.1 | QQE14280.1 | USA | F11L | NP → NP | -2.60 | 0.633 | 0.098 | 10.99 | 117.89 |
| var30 | MW374919.1 | QPZ55839.1 | USA | I13V | NP → NP | -0.54 | 0.629 | 0.096 | 14.08 | 105.00 |
| var31 | MW333012.1 | QPM32747.1 | USA | A8S | NP → P | -0.92 | 0.568 | 0.099 | 16.06 | 105.00 |
| var32 | MW310442.1 | QPK13615.1 | USA | Q29H | P → P | -1.61 | 0.645 | 0.092 | 15.81 | 107.63 |
| var33 | MW277047.1 | QPF60767.1 | Australia | A8T | NP → P | -1.69 | 0.517 | 0.096 | 19.31 | 105.00 |
| var34 | MW197470.1 | QOW16934.1 | USA | V32L | NP → NP | -1.27 | 0.626 | 0.097 | 16.06 | 110.26 |
| var35 | MW183960.1 | QOT47966.1 | Australia | Y3C | NP → P | 0.66 | 0.737 | 0.091 | 18.29 | 107.63 |
| var36 | MW070092.1 | QOH29638.1 | USA | F35C | NP → P | -2.55 | 0.629 | 0.096 | 19.26 | 107.63 |
| var37 | MW056151.1 | QOE87730.1 | USA | M21K | NP → P | -2.10 | 0.484 | 0.101 | 16.06 | 107.63 |
| var38 | MW030265.1 | QNV50343.1 | Peru | C19F | P → NP | -0.86 | 0.645 | 0.096 | 16.06 | 107.63 |
| var39 | MT994849.1 | QNR60413.1 | Iran | Y14H | NP → NP | -1.27 | 0.587 | 0.099 | 19.31 | 107.63 |
| var40 | MT981452.1 | QNQ16982.1 | USA | F7L | NP → NP | -2.45 | 0.663 | 0.097 | 16.06 | 117.89 |
| var41 | MT919771.1 | QNJ45359.1 | Philippines | V33I | NP → NP | -0.94 | 0.645 | 0.090 | 16.06 | 110.26 |

**Figure 5.**
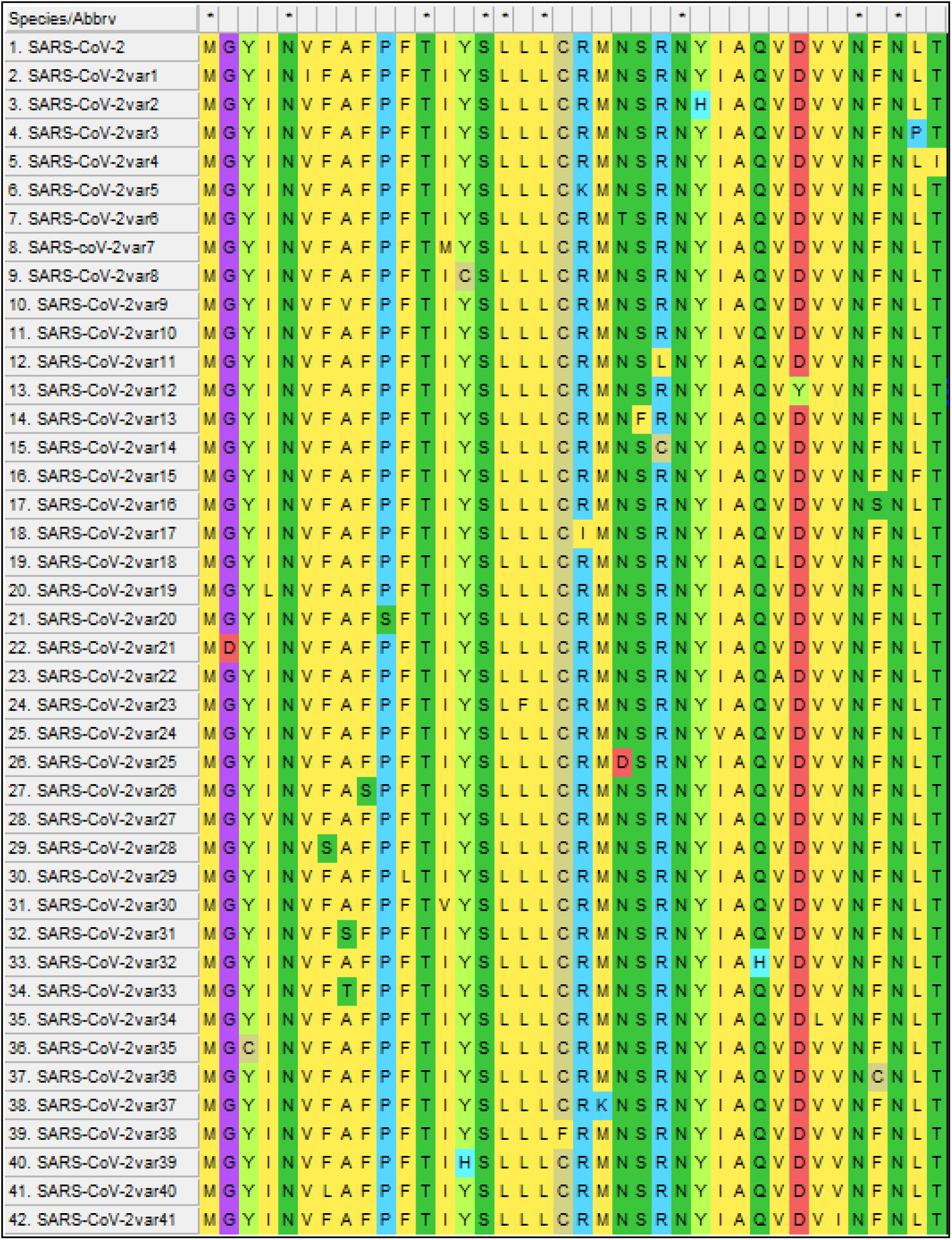
Screen capture of the MUSCLE alignment consisting of the reference ORF10 protein in SARS-CoV-2 and the forty-one variants. Conserved positions are indicated with an asterisk.

## 4. Discussion

In this study, the evolutionary history for ORF10 is similar to the history previously described by Cagliani *et al*. 2020.^14^ Currently, there exists two distinct and conserved lineages for the ORF10 in CoVs: the SARS-CoV and SARS-CoV-2 lineages. Both lineages are easily distinguishable based upon whether they encode for either a truncated (SARS-CoV lineage) or full-length (SARS-CoV-2 lineage) protein. The SARS-CoV lineage is undergoing purifying selection whereas the SARS-CoV-2 lineage is undergoing positive selection without any relaxed constraints. A bat-CoV (SARS-related-CoV Rc-o319) isolated from Japan in 2013 possessed an uninterrupted ORF10 sequence with approximately 95% nucleotide and 87% amino acid similarity to ORF10 in SARS-CoV-2.^10^ Following the addition of this new sequence, this study provides an updated phylogenetic and evolutionary look at ORF10 and continues to reinforce the current hypothesis that positive selection is occurring within the SARS-CoV-2 lineage. These data also suggest that ORF10 might encode a conserved and functional protein in SARS-CoV-2 and other closely related bat and pangolin CoVs. This lineage does have greater amino acid variations within the C-terminal; however, the cause (if any exist) behind this remains unclear.

Although the SARS-CoV lineage was not a major focus, it is worth noting that in contrast to the work performed by Cagliani *et al*. 2020 they found the SARS-CoV lineage was being subjected to neutral evolution.^14^ As such, a more detailed evolutionary analysis could help determine if the SARS-CoV lineage is under either purifying selection or neutral evolution.

The prediction of secondary structural elements for the ORF10 protein describe the presence of one α-helix and two β-strands. When compared to the structural assignment of residues in the ORF10 protein model, only 76% of residues match. One significant difference was the model contained a 3/10-helix at the C-terminus. Since this model is predicative and no experimentally derived structure is available for the ORF10 protein, this 3/10-helice’s existence along with its structural and functional role is unknown. The local scoring for residues in this structure were also lowest in comparison to the rest of the model. In order to better understand the ORF10 protein, it is recommended that an experimentally derived structure be obtained as to potentially further substantiate the predicted model generated during this study.

According to Hassan *et al*. 2020, a MoRF is thought to occur in residues 3-7.^12^ A unique feature associated with MoRFs is their ability to exist in semi-disordered (i.e. flexible) states until binding, after which they will assume a designated secondary structure.^43^ Therefore, it is possible residues 3-7 (or 3-8) function as a β-MoRF and that upon binding to a corresponding protein ligand, will assume the structure of a single β-strand. Evidence in support of this comes from the fact that residues in this region do associate with a β-strand, and have disorder scores reflecting on residues that possess moderate flexibility to accommodate changes in structure.

Protein disorder predictions show the ORF10 protein in SARS-CoV-2 is highly ordered. It is only near each terminus that residues begin to indicate some degree of flexibility. These results contradict those by Hassan *et al*. 2020.^12^ Their study indicates more flexible residues, including several disordered residues within the C-terminus, exist in the ORF10 protein.^12^ This disagreement is likely attributed to the method of prediction, as this study used multiple prediction algorithms to evaluate disorder. In doing so, per-residue disorder scores were not over estimated by a single algorithm; therefore, more conservative but reliable predictions were achieved.^26^ The results of this study indicate that mutations in the C-terminus instill changes in residue flexibility; however, these changes do not result in shifts towards disorder. Conversely, mutations in the N-terminus caused minimal differences in residue flexibility, and as noted in the C-terminus, these changes did not result in shifts towards disorder. Overall, the forty-one mutations analyzed do not pose any significant changes on the average disorder for the ORF10 protein in SARS-CoV-2.

An overwhelming number of missense mutations in the ORF10 protein decrease protein stability; however they do not result in a completely unstable protein according to the instability and aliphatic indices as determined by ProtParam. As such, these mutations are not likely to have major effects on structure or function; however, Hassan *et al*. 2020 does assert that some mutations are deleterious and may decrease the protein’s structural integrity.^12^ Modeling of these mutations could help determine if they would negatively impact the structure of the ORF10 protein.

The ORF10 protein in SARS-CoV-2 might be a membrane protein. Whether this protein is integrated within or peripheral to the membrane remains unclear, but given the extent of nonpolar residues it is reasonable to consider this protein as integrated. Furthermore, TM predictions suggest residues 3-19 span the membrane with the N-terminus exposed to the extracellular space or, if within an organelle, the cytoplasmic space. With additional evidence coming from ProFunc and the hydrophobicity plot, it appears residues 3-19 compose a TM region. Residues 28-36 could be another TM region as well; however, TMPred did not implicate these residues. Nonetheless, scores for these residues in the hydrophobicity plot were lower but still above the cutoff threshold.

In particular, CoVs encode multiple viroporins.^44^ For example, in SARS-CoV the 8a protein oligomerizes to form ion channels inside mitochondrial membranes that induce apoptosis by depolarizing membrane potential.^14,44–46^ The ORF10 protein in SARS-CoV-2 may oligomerize in such a way that polar and electrically charged residues coat the inside of a pore that facilitates the transport of ions or small molecules for viral replication, virulence, and/or pathogenicity. Based on the ORF10 protein model, it consists almost entirely of one β-α-β motif. These motifs are usually found within proteins that associate with porins. A prior study found the ORF10 protein colocalized with accessory proteins 7b and 8 in the endoplasmic reticulum (ER); however, its role in the ER was not described.^47^ In this case, the ORF10 protein may oligomerize and form a viroporin that associates with ER membranes as a means to assist in viral replication and/or processing of other viral proteins. Alternatively, the ORF10 protein could be interacting with another protein. As previously stated, the ORF10 protein was suggested to associate with the substrate adapter ZYG11B.^13^ This interaction is likely mediated by residues in the N-terminal half on account of the β-MoRF and high protein-binding propensity scores occurring throughout residues 1-14. It is tempting to think the ORF10 protein may be multifunctional; however, further studies are necessary to explore this idea.

A major limitation to this study is the quality of sequences due to the varying methods used to acquire them. Although unlikely, there is always the possibility of errors and missing information that may affect the results obtained herein. In addition, with only six sequences used, the evidence of positive selection should not be regarded as conclusive but as additional support for the hypothesis that positive selection is acting on the SARS-CoV-2 lineage for ORF10. Additional sequences of good quality from other closely related CoVs can help to resolve this matter. It is also important to note that mutations analyzed in this study are likely present in other geographic locations; therefore, the list of locations in this study should not be viewed as exhaustive.

Overexpression of ORF10 has been shown to occur in severe cases of COVID-19 whereas in milder cases its expression seems to be minimal.^16^ Therefore, knowing the structure and biochemical characteristics associated with the ORF10 protein might aid in developing diagnostic tests that detect for the ORF10 protein, thereby helping to determine the likely progression the disease may take in patients.^16^ In addition, based on results by DoGSiteScorer the ORF10 protein would not make for a suitable drug target. Despite not serving as a likely drug target, the predicted structure for the ORF10 protein may be useful in further computational analyses detailing events in viral pathogenesis or virulence. Hopefully, this study can also provide a foundation that drives further work assessing the biological relevance of ORF10 and its putatively encoded protein in SARS-CoV-2, thereby aiding in diagnostic and possibly vaccine development.

## Acknowledgments

I would like to thank the previous researchers who had deposited the sequences used in this study into NCBI. Without access to those sequences, the research described in this manuscript could have never been accomplished.

## Conflict of Interest

The author declares that no conflicts of interest exist.

## Funding Information

The work described in this article received no specific grant from any funding agency.

## Notes

### Competing Interest Statement

The authors have declared no competing interest.

## References

1 WHO: World Health Organization. (2020). “Weekly Epidemiological Update Coronavirus disease 2019 (COVID-19) 2 February 2021.” Coronavirus disease (COVID-19) Weekly Epidemiological Update and Weekly Operational Update. World Health Organization. https://www.who.int/emergencies/diseases/novel-coronavirus-2019/situation-reports

2 Wu A, Peng Y, Huang B, et al. (2020) “Genome Composition and Divergence of the Novel Coronavirus (2019-nCoV) Originating in China.” Cell Host & Microbe. 27. 3. Pg 325–328. https://doi.org/10.1016/j.chom.2020.02.001

3 Kim D, Lee JY, Yang JS, et al. (2020). “The Architecture of SARS-CoV-2 Transcriptome.” Cell. 181. Pg 914–921. https://doi.org/10.1016/j.cell.2020.04.011

4 Narayanan K, Huang C, & Makino S. (2008). “SARS coronavirus accessory proteins.” Virus Research. 133. 1. Pg 113–121. https://doi.org/10.1016/j.virusres.2007.10.009

5 Ren Y, Shu T, Wu D, et al. (2020). “The ORF3a protein of SARS-CoV-2 induces apoptosis in cells.” Cell and Molecular Immunology. 17. Pg 881–883. https://doi.org/10.1038/s41423-020-0485-9

6 Baruah C, Devi P, & Sharma DK. (2020). “Sequence Analysis and Structure Prediction of SARS-CoV-2 Accessory Proteins 9b and ORF14: Evolutionary Analysis Indicates Close Relatedness to Bat Coronavirus.” BioMed Research International. 7234961. Pg 1–13. https://doi.org/10.1155/2020/7234961

7 George T, Rawlinson D, Featherstone L, et al. (2020). “Direct RNA sequencing and early evolution of SARS-CoV-2.” bioRxiv. Preprint. https://doi.org/10.1101/2020.03.05.976167

8 Pancer K, Milewska A, Owczarek K, et al. (2020). “The SARS-CoV-2 ORF10 is not essential in vitro or in vivo in humans.” PLoS Pathogens. 16. 12. e1008959. https://doi.org/10.1371/journal.ppat.1008959

9 Khailany RA, Safdar M, and Ozaslan M. (2020). “Genomic characterization of a novel SARS-CoV-2.” Gene Reports. 19. 100682. https://doi.org/10.1016/j.genrep.2020.100682

10 Murakami S, Kitamura T, Suzuki J, et al. (2020). “Detection and Characterization of Bat Sarbecovirus Phylogenetically Related to SARS-CoV-2, Japan.” Emerging Infectious Diseases. 26. 12. Pg. 3025–3029. https://dx.doi.org/10.3201/eid2612.203386.

11 Liu P, Jiang JZ, Wan XF, et al. (2020) “Are pangolins the intermediate host of the 2019 novel coronavirus (SARS-CoV-2)?” PLoS Pathogens. 16. 5. Pg 1–13. https://doi.org/10.1371/journal.ppat.1008421

12 Hassan S, Attrish D, Ghosh S, et al. (2020). “Notable sequence homology of the ORF10 protein introspects the architecture of SARS-COV-2.” bioRxiv. Preprint. https://doi.org/10.1101/2020.09.06.284976

13 Gordon DE, Jang GM, Bouhaddou M, et al. (2020). “A SARS-CoV-2 protein interaction map reveals targets for drug repurposing.” Nature. 583. Pg 459–468. https://doi.org/10.1038/s41586-020-2286-9

14 Cagliani R, Forni D, Clerici M, et al. (2020). “Coding potential and sequence conservation of SARS-CoV-2 and related animal viruses.” Infection, Genetics, and Evolution. 83. 104353. https://doi.org/10.1016/j.meegid.2020.104353

15 Mishra S. (2020). “Designing of cytotoxic and helper T cell epitope map provides insights into the highly contagious nature of the pandemic novel coronavirus SARS-CoV-2.” Royal Society Open Science. 7201141. http://doi.org/10.1098/rsos.201141

16 Liu T, Jia P, Fang B, et al. (2020). “Differential Expression of Viral Transcripts From Single-Cell RNA Sequencing of Moderate and Severe COVID-19 Patients and Its Implications for Case Severity.” Frontiers in Microbiology. 11. 603509. https://doi.org/10.3389/fmicb.2020.603509

17 Michel CJ, Mayer C, Poch O, et al. (2020). “Characterization of accessory genes in coronavirus genomes.” Virology Journal. 17. 131. https://doi.org/10.1186/s12985-020-01402-1

18 Zhang Z, Schwartz S, Wagner L, et al. (2000). “A greedy algorithm for aligning DNA sequences.” Journal of Computational Biology: A Journal of Computational Molecular Cell Biology. 7. (1-2). Pg 203–214. https://doi.org/10.1089/10665270050081478

19 Edgar RC. (2004) “MUSCLE: multiple sequence alignment with high accuracy and high throughput.” Nucleic Acids Research. 32. 5. Pg. 1792–1797. https://doi.org/10.1093/nar/gkh340

20 Kumar S, Stecher G, Li M, et al. (2018) “MEGA X: Molecular Evolutionary Genetics Analysis across computing platforms.” Molecular Biology and Evolution. 35. Pg 1547–1549. https://doi.org/10.1093/molbev

21 Hall BG. (2018). “Phylogenetic Trees Made Easy: A How-To Manual.” Oxford University Press, New York. 5. Pg 60–61.

22 Saitou N and Nei M. (1987). “The neighbor-joining method: A new method for reconstructing phylogenetic trees.” Molecular Biology and Evolution. 4. Pg 406–425. https://doi.org/10.1093/oxfordjournals.molbev.a040454

23 Kosakovsky SL & Frost SD. (2005). “Not so different after all: a comparison of methods for detecting amino acid sites under selection.” Molecular Biology and Evolution. 22. 5. Pg. 1208–1222. https://doi.org/10.1093/molbev/msi105

24 Wertheim JO, Murrell B, Smith MD, et al. (2015). “RELAX: detecting relaxed selection in a phylogenetic framework.” Molecular Biology and Evolution. 32. 3. Pg. 820–832. https://doi.org/10.1093/molbev/msu400

25 Kabat EA, Wu TT, & Bilofsky H. (1977). “Unusual distribution of amino acids in complementarity-determining (hypervariable) segments of heavy and light chains of immunoglobulins and their possible roles in specificity of antibody combining sites.” Journal of Biological Chemistry. 252. Pg. 6609–6616. https://www.jbc.org/content/252/19/6609.full.pdf

26 Cleveland SB, Davies J, and McClure MA. (2011) “A Bioinformatics Approach to the Structure, Function, and Evolution of the Nucleoprotein of the Order Mononegavirales.” PLoS ONE. 6.5. Pg 1–13. https://doi.org/10.1371/journal.pone.0019275

27 Gasteiger E, Hoogland C, Gattiker A, et al. (2005). “The Proteomics Protocols Handbook.” Humana Press. Pg 571–607.

28 Zhang J & Kurgan L. (2019). “SCRIBER: accurate and partner type-specific prediction of protein-binding residues from proteins sequences.” Bioinformatics. 35.14. Pg 343–353. https://doi.org/10.1093/bioinformatics/btz324

29 Jones DT. (1999). “Protein secondary structure prediction based on position-specific scoring matrices.” Journal of Molecular Biology. 292. Pg 195–202. https://doi.org/10.1006/jmbi.1999.3091

30 Hofmann K & Stoffel W. (1993). “TMbase: A database of membrane spanning proteins segments.” Biological Chemistry Hoppe-Seyler. 374. 166.

31 McGuffin LJ, Adiyaman R, Maghrabi AHA, et al. (2019). “IntFOLD: an integrated web resource for high performance protein structure and function prediction.” Nucleic Acids Research. 47. Pg 408–413. https://doi.org/10.1093/nar/gkz322

32 Bhattacharya D & Cheng J. (2012). “3Drefine: Consistent Protein Structure Refinement by Optimizing Hydrogen Bonding Network and Atomic Level Energy Minimization.” Proteins: Structure, Function, and Bioinformatics. 81.1. Pg 119–131. https://doi.org/10.1002/prot.24167

33 Benkert P, Biasini M, & Schwede T. (2012). “Toward the estimation of the absolute quality of individual protein structure models.” Bioinformatics. 27. Pg 343–350. https://doi.org/10.1093/bioinformatics/btq662

34 Colovos C and Yeates TO. (1993) “Verification of protein structures: patterns of nonbonded atomic interactions.” Protein Science. 2. 9. Pg 1511–1519. https://doi.org/10.1002/pro.5560020916

35 Laskowski RA, Rullmann JAC, MacArthur MW, et al. (1996) “AQUA and PROCHECK-NMR: Programs for checking the quality of protein structures solved by NMR.” Journal of Biomolecular NMR. 8. Pg 477–486. https://doi.org/10.1007/BF00228148

36 Laskowski RA, MacArthur MW, Moss DS, et al. (1993) “PROCHECK: a program to check the stereochemical quality of protein structures.” Journal of Applied Crystallography. 26. Pg 283–291. https://doi.org/10.1107/S0021889892009944

37 Mirsha S. (2020). “ORF10: Molecular Insights into the Contagious Nature of Pandemic Novel Coronavirus 2019-nCoV.” ChemRxiv. Preprint. https://doi.org/10.26434/chemrxiv.12118839.v3

38 Zhang Lab. University of Michigan. zhanglab.ccmb.med.umich.edu/COVID-19/. Accessed 15 Sept. 2020.

39 Roman LA, James WD, & Janet MT. (2005). “ProFunc: a server for predicting protein function from 3D structure.” Nucleic Acids Research. 33. Pg. W89–W93. https://doi.org/10.1093/nar/gki414

40 Andrea V, Daniel K, Friedrich R, et al. (2012). “DoGSiteScorer: a web server for automatic binding site prediction, analysis and druggability assessment.” Bioinformatics. 28. 15. Pg 2074–2075. https://doi.org/10.1093/bioinformatics/bts310

41 Pettersen EF, Goddard TD, Huang CC, et al. (2004) “UCSF Chimera-a visualization system for exploratory research and analysis.” Journal of Computational Chemistry. 25.13. Pg 1605–1612. https://doi.org/10.1002/jcc.20084

42 Emidio C, Piero F, & Rita C. (2005). “I-Mutant2.0: predicting stability changes upon mutation from the protein sequence or structure.” Nucleic Acids Research. 33. Pg. W306–W310, https://doi.org/10.1093/nar/gki375

43 Mohan A, Oldfield CJ, Radivojac P, et al. (2006). “Analysis of molecular recognition features (MoRFs).” Journal of Molecular Biology. 362.5. Pg 1043–1059. https://doi.org/10.1016/j.jmb.2006.07.087

44 Forni D, Cagliani R, Clerici M, et al. (2017). “Molecular Evolution of Human Coronavirus Genomes.” Trends in Microbiology. 25.1. Pg 35–48. https://doi.org/10.1016/j.tim.2016.09.001

45 Chen CC, Krüger J, Sramala I, et al. (2011). “ORF8a of SARS-CoV forms an ion channel: Experiments and molecular dynamics simulations.” Biochimica et Biophysica Acta (BBA) - Biomembranes. 1808.2. Pg 572–579. https://doi.org/10.1016/j.bbamem.2010.08.004

46 Chen CY, Ping YH, Lee HC, et al. (2007). “Open reading frame 8a of the human severe acute respiratory syndrome coronavirus not only promotes viral replication but also induces apoptosis.” The Journal of Infectious Diseases. 196.3. Pg 405–415. https://doi.org/10.1086/519166

47 Zhang J, Cruz-cosme R, Zhuang MW, et al. (2020). “A Systemic and Molecular Study of Subcellular Localization of SARS-CoV-2 Proteins.” Signal Transduction and Targeted Therapy. 5. 269. https://doi.org/10.1038/s41392-020-00372-8

