## Supplementary Data for "Characterization and structural prediction of the putative ORF10 protein in SARS-CoV-2"

| **CoV Species** | **Nucleotide Accession** | **Nucleotide Similarity (%)** |
| --- | --- | --- |
| Bat SARS-CoV Rp3 | DQ071615.1 | 90 |
| Bat SARS-like-CoV Rf4092 | KY417145.1 | 91 |
| BtRf-BetaCoV/JL2012 | KJ473811.1 | 92 |
| Bat SARS-like CoV Rs4231 | KY417146.1 | 92 |
| Bat SARS-like CoV As6526 | KY417142.1 | 92 |
| SARS-related Bat-CoV Longquan-140 | KF294457.1 | 92 |
| Bat-SARS-CoV HKU3-1 | DQ022305.2 | 92 |
| Rhinolophus affinis CoV LYRa11 | KF569996.1 | 93 |
| BtRs-BetaCoV/HuB2013 | KJ473814.1 | 93 |
| Bat-CoV Cp/Yunnan2011 | JX993988.1 | 93 |
| Bat-CoV Rp/Shaanxi2011 | JX993987.1 | 93 |
| SARS-CoV Tor2 | JX163928.1 | 93 |
| Bat SARS-CoV Rm1 | DQ412043.1 | 93 |
| Bat-CoV BM48-31/BGR/2008 | GU190215.1 | 93 |
| SARS-related CoV BtKY72 | KY352407.1 | 94 |
| Pangolin-CoV CoV GX-P1E | MT040334.1 | 94 |
| SARS-related-CoV Rc-o319 | LC556375.1 | 95 |
| Pangolin-CoV MP789 | MT121216.1 | 99 |
| bat-SL-CoVZC45 | MG772933.1 | 99 |
| Bat-CoV RaTG13 | MN996532.2 | 99 |

**Table S1.** List of CoV species used in this study. The nucleotide similarity corresponds to the ORF10 sequences in each CoV when compared against the ORF10 sequence in SARS-CoV-2.

| **Position** | **Residue** | **Depp Score** |
| --- | --- | --- |
| 3 | Y | 0.0657 |
| 12 | T | 0.0783 |
| 14 | Y | 0.0123 |
| 15 | S | 0.0376 |
| 23 | S | 0.1393 |
| 26 | Y | 0.0445 |
| 38 | T | 0.0027 |

**Table S2.** Prediction scores for phosphorylation sites in the ORF10 protein. Scores ≥ 0.5 indicate a higher likelihood of being phosphorylated. However, none of the residues listed in this table were found to be a potential phosphorylation site in the SARS-CoV-2 ORF10 protein.

| **Protein Data** | | **PSIPRED v4.0** | | | **Structural Assignment** |
| --- | --- | --- | --- | --- | --- |
| **Position** | **Residue** | **H** | **E** | **C** |  |
| 1 | M | 0.001 | 0.002 | 0.999 | C |
| 2 | G | 0.024 | 0.095 | 0.885 | C |
| 3 | Y | 0.029 | 0.388 | 0.564 | C |
| 4 | I | 0.014 | 0.597 | 0.353 | E |
| 5 | N | 0.015 | 0.755 | 0.204 | E |
| 6 | V | 0.132 | 0.495 | 0.335 | E |
| 7 | F | 0.155 | 0.503 | 0.345 | E |
| 8 | A | 0.081 | 0.645 | 0.265 | E |
| 9 | F | 0.123 | 0.412 | 0.499 | C |
| 10 | P | 0.165 | 0.134 | 0.735 | C |
| 11 | F | 0.695 | 0.054 | 0.289 | H |
| 12 | T | 0.789 | 0.113 | 0.154 | H |
| 13 | I | 0.939 | 0.057 | 0.055 | H |
| 14 | Y | 0.989 | 0.012 | 0.009 | H |
| 15 | S | 0.984 | 0.005 | 0.015 | H |
| 16 | L | 0.986 | 0.005 | 0.013 | H |
| 17 | L | 0.970 | 0.010 | 0.030 | H |
| 18 | L | 0.932 | 0.036 | 0.070 | H |
| 19 | C | 0.865 | 0.061 | 0.110 | H |
| 20 | R | 0.861 | 0.050 | 0.134 | H |
| 21 | M | 0.717 | 0.016 | 0.262 | H |
| 22 | N | 0.428 | 0.015 | 0.571 | C |
| 23 | S | 0.084 | 0.018 | 0.912 | C |
| 24 | R | 0.021 | 0.031 | 0.956 | C |
| 25 | N | 0.052 | 0.144 | 0.780 | C |
| 26 | Y | 0.020 | 0.505 | 0.429 | E |
| 27 | I | 0.022 | 0.821 | 0.133 | E |
| 28 | A | 0.005 | 0.931 | 0.057 | E |
| 29 | Q | 0.005 | 0.908 | 0.080 | E |
| 30 | V | 0.006 | 0.835 | 0.114 | E |
| 31 | D | 0.008 | 0.713 | 0.232 | E |
| 32 | V | 0.015 | 0.756 | 0.138 | E |
| 33 | V | 0.006 | 0.916 | 0.071 | E |
| 34 | N | 0.010 | 0.845 | 0.127 | E |
| 35 | F | 0.008 | 0.497 | 0.538 | C |
| 36 | N | 0.004 | 0.604 | 0.391 | E |
| 37 | L | 0.008 | 0.124 | 0.864 | C |
| 38 | T | 0.000 | 0.001 | 0.999 | C |

**Table S3.** Per-residue secondary structure prediction scores provided by PSIPRED outlining the probability of finding a residue in either a helix (H), strand (E), or coil (C). The highest of the three scores indicated the structural designation of that specific residue.

| **SARS-CoV-2 ORF10 Protein Models** | | |
| --- | --- | --- |
| **Model Type** | **p-value** | **Model Quality** |
| Model 1 | 3.33E-03 | 0.3714 |
| Model 2 | 1.06E-02 | 0.3577 |
| Model 3 | 1.25E-02 | 0.3558 |
| Model 4 | 4.67E-02 | 0.3378 |
| Model 5 | 5.54E-02 | 0.3343 |

**Table S4.** Quality and confidence (p-value) scores corresponding to all five generated ORF10 protein models using the IntFOLD web server.

| **Model ID** | **QMEAN** |
| --- | --- |
| Model 1 | -0.88 |
| Model 2 | -0.68 |
| Model 3 | -0.81 |
| Model 4 | -0.66 |
| Model 5 | -0.70 |

**Table S5.** QMEAN global Z-scores corresponding to all five refined ORF10 protein models using the 3Drefine web server.

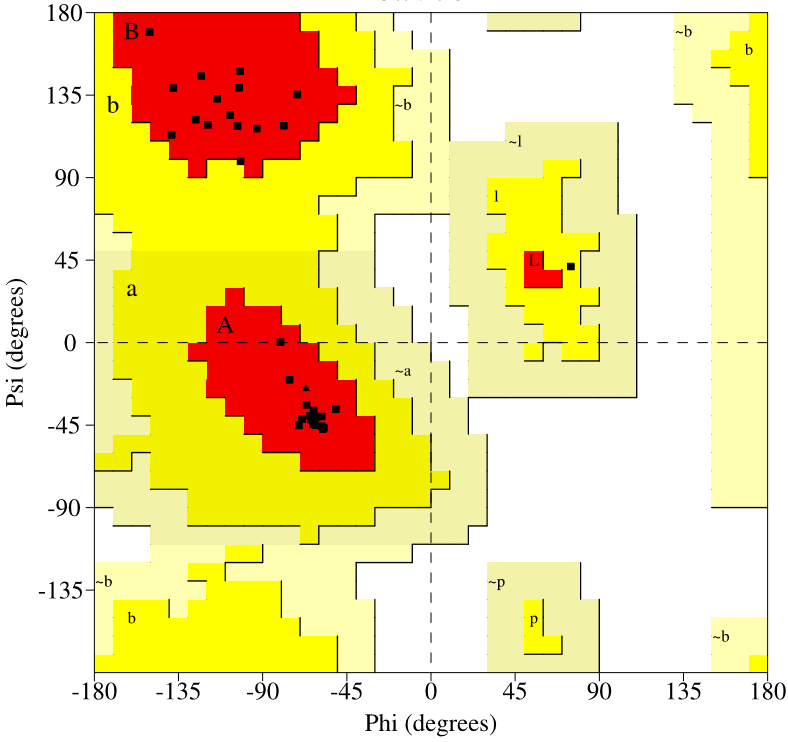

**Figure S1. (A)** Ramachandran plot for the predicted ORF10 protein model. [A,B,L] = favoured region. [a,b,l,p] = allowed regions. [~a,~b,~l,~p] = generously allowed regions. All of the glycine residues are shown as triangles.

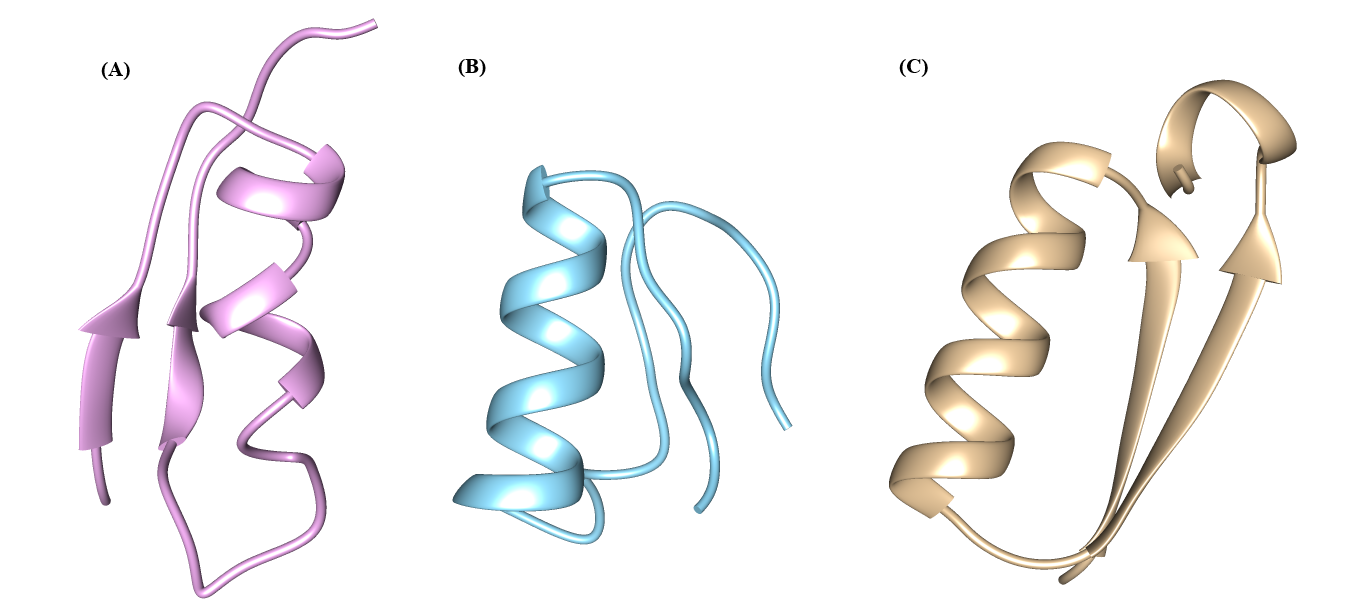

**Figure S2.** Three ribbon diagrams generated by different methods to represent the ORF10 protein. The **(A)** I-TASSER, **(B)** QUARK, and (**C)** current study's model of the ORF10 protein. For clarification, the colors of each model are not relevant other than helping to further distinguish between models.

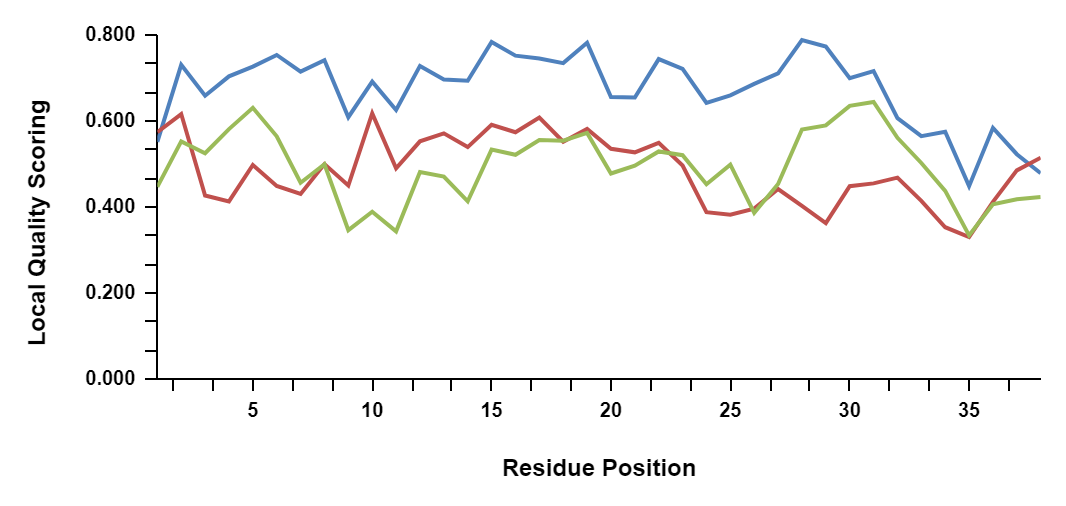

**Figure S3.** Local quality scoring for each ORF10 protein model. Comparisons were performed against three different models: I-TASSER (green), QUARK (red), and this current study’s model (blue) of the ORF10 protein.
